# Early maltreatment experience increases resting-state functional connectivity in the theory of mind (ToM) network

**DOI:** 10.1101/493635

**Authors:** Sara Boccadoro, Roma Siugzdaite, Anna R. Hudson, Lien Maeyens, Charlotte Van Hamme, Sven C. Mueller

## Abstract

**Background:** Early life stressful events, such as childhood maltreatment, significantly increase risk for the development of psychopathology and are associated with impairments in socio-cognitive skills including theory-of-mind (ToM). However, to date, no study has examined the resting-state activity of the ToM network in adults with maltreatment history.

**Methods:** Thirty-five women with a history of childhood maltreatment and 31 unaffected women completed a resting-state scan and a ToM localizer task. The peak coordinates from the localizer were used as the seed regions for the resting-state functional connectivity (RSFC) analyses (temporoparietal junction, dorsomedial prefrontal cortex, middle temporal gyrus and precuneus).

**Results:** Child abuse was associated with increased RSFC between various ToM regions including the precuneus and the brainstem suggesting altered hierarchical processing in ToM regions. In addition, RSFC was also changed between the dorsomedial prefrontal cortex and the cerebellum.

**Limitations:** A small and heterogeneous sample.

**Conclusions:** The data indicate a lasting influence of early life stress on neural networks involved in social processing and may underlie the social difficulties reported by maltreated individuals.

## Introduction

Childhood maltreatment is a major public health issue. According to official estimates, around 35% of adult women in the European Union have experienced physical, psychological, or sexual violence before the age of 15 (1). Early life stressful (ELS) experiences such as sexual, physical, and emotional abuse can lead to the development of behavior problems and affect neural structure and plasticity. These then result in a predisposition towards the development of mental disorders in adulthood, including mood and anxiety disorders, post-traumatic stress disorder (PTSD), substance abuse and personality disorders (2, 3). One important and widely accepted consequence of any type of childhood maltreatment is an impairment in socio-emotional development, leading to poorer performance in critical functions such as emotion recognition and understanding, and theory-of-mind abilities (4).

Because of the inherent violation of personal integrity and autonomy, maltreated children may have a different perception of interpersonal interactions and exhibit changes in social abilities, emotions and other mental states associated with such interactions, than children without maltreatment experience (4). Social understanding then is the ability to interpret the thoughts and feelings of other people, the so-called “theory of mind” (ToM). Specifically, ToM is the core human socio-cognitive ability to explain and predict the behavior of others by attributing to them independent mental states, such as beliefs, emotions, intentions, hopes and desires that may be different from our own (5).

Such complex functions recruit a diverse array of brain areas that (presumably) synchronize with one another to integrate different pieces of information to represent ToM at the neural level (e.g., areas involved in self-other distinction, self-referential processing, episodic memory, attention, language)(6). According to recent reviews and meta-analyses (6, 7) neural nodes belonging to this ToM network consist of: 1) the *temporo-parietal junction* (TPJ) (Brodmann Area 39), 2) the *precuneus* (mesial extent of Brodmann’s Area 7), 3) the *dorsomedial prefrontal cortex (dmPFC)*, and 4) the *middle temporal gyrus* (MTG, BA 21). However, given that ELS may alter specific aspects of ToM (4), one would hypothesize a perturbation of the basic neural oscillatory network involved in ToM after such experiences.

Characterization of resting-state brain networks in persons with mental illness or at-risk may help to identify perturbations of underlying neural oscillations and changed brain dynamics. Within trauma research, only a few studies to date have examined intrinsic functional connectivity (FC) but with much heterogeneity in findings (8–12). Decreased FC between the amygdala and other brain areas such as the putamen, insula, or subgenual anterior cingulate cortex have been documented in maltreated adults (10) and children (8). Directly contrasting these findings, a task-based study reported increased (rather than decreased) FC for traumatized participants relative to comparisons specifically between midbrain structures (superior colliculus (SC) and locus coeruleus (LC)) and subcortical brain areas including the amygdala, the anterior cingulate, the thalamus, and the striatum (12). In their (12) study, participants viewed animated video sequences of faces with varying emotional expressions and direct or averted gaze. The authors argued that this increased FC may reflect preferential recruitment of a fast subcortical processing route indicative of an innate threat system that is quickly (and probably standardly) active in traumatized persons. Indeed, a hallmark symptom of PTSD is a constant level of alert (hypervigilance), an issue that clinicians aim to moderate before commencing other therapy (13). However, the findings by Steuwe et al. (12) have to be considered tentative as they were 1) task-based and consisted of a re-analysis of a prior fMRI study (14), and 2) were comprised of a relatively small sample (N=16 per group).

Given these discrepancies and the interesting hypothesis that an innate state of fear after ELS may influence social processing, this study assessed the critical question to what extent individuals with and without ELS show altered resting-state functional connectivity (RSFC) in a ToM network including structures relevant to processing of social signals. To this end, well-known ToM regions were identified in the present sample by virtue of an independent localizer task. If Steuwe et al. are correct and low level fast processing interferes with higher order socio-affective function, then we would also expect to see increased FC between these ToM regions (and possibly the midbrain). Such evidence would then demonstrate long-term neural oscillatory changes after ELS in a social processing network. Importantly, such identification would aid in the continued quest for neuroimaging biomarkers of stress-related disorders (15).

## Methods

### Participants

Thirty-five women with a history of childhood sexual, physical, or emotional abuse (CA) and 31 comparison women without such history or history of other, non-childhood abuse related traumas in either childhood or adulthood (UC) participated (Table 1). Participants did not differ in age, education, or medication usage but CA participants, relative to UC, had significantly higher levels of depression (t(49.48) = 5.03, *p* < 0.001), dissociative experiences (t(45.043) = 3.535, *p* = 0.001), current psychopathology (χ^2^(1) = 26.615, *p* < 0.001), trait anxiety (t(63) = 5.63, *p* < 0.001) and state anxiety (t(64) = 4.55, *p* < 0.001). However, UC had higher resilience than CA (t(64) = −3.092, *p* = 0.003). In addition, CA participants also had significantly higher levels of self-reported empathy (t(64) = 3.05, *p* = 0.003).

**Table 1.** Demographic and clinical characteristics of the child abuse (CA) and the unaffected comparisons (UC)

|  | Child abuse group<br>(CA)<br>N = 35 | Unaffected comparisons<br>(UC)<br>N = 31 | *p* |
| --- | --- | --- | --- |
| Age (years), mean (SD) | 36.16 (12.19) | 36.42 (11.46) | 0.929 |
| Highest education level (n)<sup>a</sup> |  |  |  |
| High school | 10 | 6 | 0.397 |
| Bachelor | 16 | 13 | 0.174 |
| Master/PhD/Postdoc | 4 | 7 | 0.451 |
| Psychiatric Medication (n) | 7 | 2 | 0.109 |
| Depression (BDI) score (SD) | 15.68 (10.61) | 5.35 (5.30) | <0.001 |
| State anxiety (STAI) score (SD) | 39.17 (8.22) | 30.45 (7.22) | <0.001 |
| Trait anxiety (STAI) score (SD) | 47.81 (8.81) | 35.19 (9.24) | <0.001 |
| Resilience (RS) score (SD) | 126.86 (17.10) | 138.61 (13.25) | 0.001 |
| Dissociative Experience (DES) score (SD) | 20.50 (12.99) | 10.12 (7.73) | 0.001 |
| Self-reported empathy (IRI) score (SD) | 72.8 (12.43) | 64.48 (9.29) | 0.001 |
| IRI – Perspective taking (SD) | 18.89 (4.81) | 18.03 (3.99) | 0.439 |
| IRI – Fantasy (SD) | 17.89 (5.51) | 16.13 (5.64) | 0.200 |
| IRI – Empathic concern (SD) | 21.14 (4.39) | 20.10 (4.22) | 0.329 |
| IRI – Personal distress (SD) | 14.86 (4.17) | 10.23 (4.46) | <0.001 |
| Current psychopathology (at least one) | 27 | 6 | <0.001 |
| Mood Disorders (n) | 13 | 2 | 0.001 |
| Anxiety Disorders (n) | 19 | 2 | <0.001 |
| PTSD (n) | 7 | 0 | 0.008 |
| Alcohol dependence and abuse (n) | 0 | 2 | 0.127 |
| Substance dependence and abuse (n) | 0 | 2 | 0.127 |
| Psychotic Disorders (n) | 1 | 0 | 0.351 |
| Eating Disorders (n) | 2 | 1 | 0.528 |
| Personality Disorders (n)<sup>d</sup> | 13 | 0 | <0.001 |
| Suicidality | 10 | 0 | 0.001 |
| Type of abuse |  |  |  |

Participants were recruited via self-help groups (CA only), flyers, and social media. Participants were matched for age, sex, handedness, and level of education. The study was approved by the ethical committee of Ghent University Hospital and all participants provided written informed consent prior to the study and received 30 EUR compensation.

### ToM localizer

A Dutch translation of a previously validated ToM localizer (~10 mins) was used (16) to identify brain regions involved in ToM, which were then used for seed-based functional connectivity (FC) analysis. Participants read 20 short stories presented in fixed order pertaining either to: characters and their false beliefs about the world (“false belief’ condition), or to inanimate objects such as maps or photographs, which displayed false information (“false photograph” condition). Half of the stories (10) belonged to the “false belief’ condition whereas the other half (10) belonged to the “false photograph” condition. Each story was presented for 10,000 ms. Participants then read a statement, presented for 10,000 ms, on that story and responded using an MRI compatible response box (Cedrus)(index finger = true statement, middle finger = false statement). The story and statement phases were conflated and analyzed together. First-level models contained separate regressors for each condition (“false belief’ and “false photograph”) as well as subject movement parameters. To identify regions involved in ToM, a one-sample t-test using the whole group was performed on the false belief > false photograph contrast at the whole brain level, with age, depression scores and trait anxiety scores as regressors of no interest. The coordinates of peak activation (corrected for multiple comparisons (family-wise error, FWE) and thresholded at *p* < .05 and cluster size *k* > 10) were then used in the seed-based resting-state FC analyses. There were no significant group differences in the localizer.

### Questionnaires

#### SLESQ

The Stressful Life Events Screening Questionnaire (SLESQ (17)) was used to assess previous traumatic exposure. Participants in the present study were classified as CA if they positively answered items 5 (rape), 6 (sexual assault), which together were considered as sexual abuse, 7 (childhood physical abuse), and/or 9 (emotional abuse), and if their age at onset of the abuse was below 17 years. The cut-off of 17 years was chosen to keep similarity with other questionnaires assessing childhood abuse, such as the Childhood Trauma Questionnaire (18). Cronbach’s alpha present sample was 0.828.

#### IRI

The 28-item Interpersonal Reactivity Index (IRI (19, 20)) measures empathic responsiveness (present α = 0.792) on 4 scales: two cognitive (perspective-taking, α = 0.735; fantasy, α = 0.797) and two affective (empathic concern, α = 0.752; personal distress, α = 0.730).

#### BDI-II

The 21-item Beck Depression Inventory-II (BDI-II (21)) measures depressive symptoms in adults according to DSM-IV criteria covering cognitive, affective, and somatic aspects of depression (present α = 0.934).

#### STAI

The State-Trait Anxiety Inventory (STAI (22, 23)) was used to measure state (20 items) and trait anxiety (20 items) (present α state: 0.869; trait: 0.604).

#### DES

The 28-item Dissociative Experiences Scale (DES (24, 25)) measures the extent to which respondents experience dissociative symptoms such as depersonalisation, derealisation, and disturbances in memory and identity in their daily life (present α = 0.897).

#### RS

The Resilience Scale (RS (26)) measures mental resilience using 26 statements about the self that are judged from “strongly disagree” to “strongly agree” on a 7-point Likert scale (present α = 0.850).

#### MINI + BPD screening

The Mini International Neuropsychiatric Interview (MINI (27, 28)) was administered by trained clinical psychology masters students. The MINI is a structured interview, assessing current and lifetime histories of Axis I disorders plus one Axis II disorder, antisocial personality disorder, based on DSM-IV criteria. Since previous research underlines a link between experience of childhood abuse and borderline personality disorder (BPD)(3), an additional MINI-style subsection assessing BPD symptomatology was included. Items for this subsection were designed based on DSM-IV criteria, translated into Dutch, and back-translated.

### Imaging data acquisition

Images were acquired as part of a larger study at Ghent University Hospital using a 3T Magnetom Siemens TrioTim MRI scanner. Firstly, a T1-weighted high resolution anatomical scan was acquired (repetition time [TR] = 2250 ms, echo time [TE] = 4.18 ms, image matrix = 256 × 256, field of view [FOV] = 256 mm, flip angle = 9°, slice thickness = 1 mm, voxel size = 1.0 × 1.0 × 1.0 mm, number of slices = 176). Then the ToM localizer scan was acquired using a T2*-weighted Echo Planar Images (EPI) sequence (TR = 2000 ms, TE = 28 ms, image matrix = 384 × 384, FOV = 224 mm, flip angle = 80°, slice thickness = 3.0 mm, voxel size = 3.5 × 3.5 × 3.0 mm, number of slices = 34). Finally, resting-state fMRI data were acquired also using a T2*-weighted Echo Planar Images (EPI) sequence (TR = 2000 ms, TE = 27 ms, image matrix = 384 × 384, FOV = 192 mm, flip angle = 90°, slice thickness = 3.0 mm, voxel size = 3.5 × 3.5 × 3.0 mm, number of slices = 34). The scanning time for resting-state data was 6:06 m.

### Image preprocessing

Resting-state fMRI data were preprocessed using SPM8 (http://www.fil.ion.ucl.ac.uk/spm/software/spm8/) on Matlab. The first 10 volumes were removed for each subject to account for signal saturation effects and the data images were preprocessed by applying slice timing, realignment to adjust for head movement during data acquisition, normalization with the standard EPI template from the Montreal Neurological Institute (MNI) (resampling voxel size = 3 × 3 × 3 mm^3^) and spatial smoothing to reduce the effects of the bad normalization by using an 8-mm full-width at half-maximum (FWHM) Gaussian kernel. The images were visually inspected to verify the validity of the normalization step for all the data.

The images were segmented (modulated) into non-brain and brain tissue (grey matter, GM; white matter, WM; cerebrospinal fluid, CSF). The functional data were then detrended and filtered with a standard low-pass filter (0.01-0.08 Hz), because only frequencies below 0.1 Hz contribute to regionally specific BOLD correlations, while faster frequencies are related to cardiac or respiratory factors (29).

Due to the fact that spontaneous activity is contaminated by artifacts such as scanner instability or non-neuronal physiological fluctuations, a regression of nuisance step was performed by regressing motion (using a brain mask), white matter (WM mask) and cerebral spinal fluid (CSF mask). All masks were created using SPM8 and visually inspected afterwards.

The last preprocessing step took care of motion using frame-wise displacement (FWD), a method developed by Power and colleagues to adjust for small head movements that may add spurious noisy variance. Their (30, 31) method measures how much the head changes position from one frame to the next and is calculated as the sum of the absolute values of the differentiated realignment estimates at every timepoint. The analysis was conducted using their BRAMILA script for framewise displacement with a recommended threshold of < 0.5 mm. Since individual subject value means were all well below 0.5, no participant was excluded (mean total FWD = 0.078 mm, FWD range = 0.033-0.217 mm). There were no significant group differences in movement parameters (FWD values).

### Seed-based resting-state functional connectivity analysis

After preprocessing, the data were analyzed using seed-based FC taking the following peak coordinates from the ToM localizer as seeds: Right TPJ (MNI coordinates xyz: 46 −58 20), Left TPJ (−48 −56 24), Precuneus (2 −56 34), dmPFC (−10 54 28), Left MTG (−56 −2 −18) and Right MTG (52 −8 −16).

The seed-based FC analysis was conducted using REST toolbox, using a user-defined mask (the brain mask) and by creating a 6-mm radius sphere around the single voxel seed. The mean activation of every seed region was calculated and then correlated to all the voxels of the brain, to look at the correlations between each time-series. Correlations were normalized using Fisher transformation, to ensure that all subjects were in the same normalized space.

Second level analysis was performed using SPM8, by entering participants’ connectivity maps with age of participants, mean FWD values and depression scores as covariates, to evaluate possible differences between CA and UC. To compare FC between CA and UC a two-sample *t*-test was performed. Given FC analyses of the seed region were examined across the whole brain, we used the updated AFNI’s 3dClustSim for correction for multiple comparisons (https://afni.nimh.nih.gov/pub/dist/doc/program_help/3dClustSim.html) with a voxelwise threshold at *p*<0.001 and the clusterwise threshold at *p*<0.01. Simulations resulted in a cluster size of at least 37 contiguous voxels.

To control for the influence of depressive symptoms, anxiety, number and severity of abuse, empathy, behavioral performance during the ToM task (accuracy) or symptoms of dissociation on the main results, a second-level analysis or correlations of scores with ROIs betas was conducted separately for controls and trauma groups. Connectivity maps of the participants of each group and the respective scores of one group at a time as a covariate were entered using the same threshold as above. Additionally, to assess influence of a PTSD diagnosis, analyses were rerun excluding all women (N=7) who would qualify for such a diagnosis. Neither depressive symptoms, anxiety, empathy, or dissociative experiences influenced the results and will not be discussed further. Importantly, diagnosis of PTSD did also not influence the results (cf. supplementary Tables, supplementary Figure 1).

**Figure 1.**
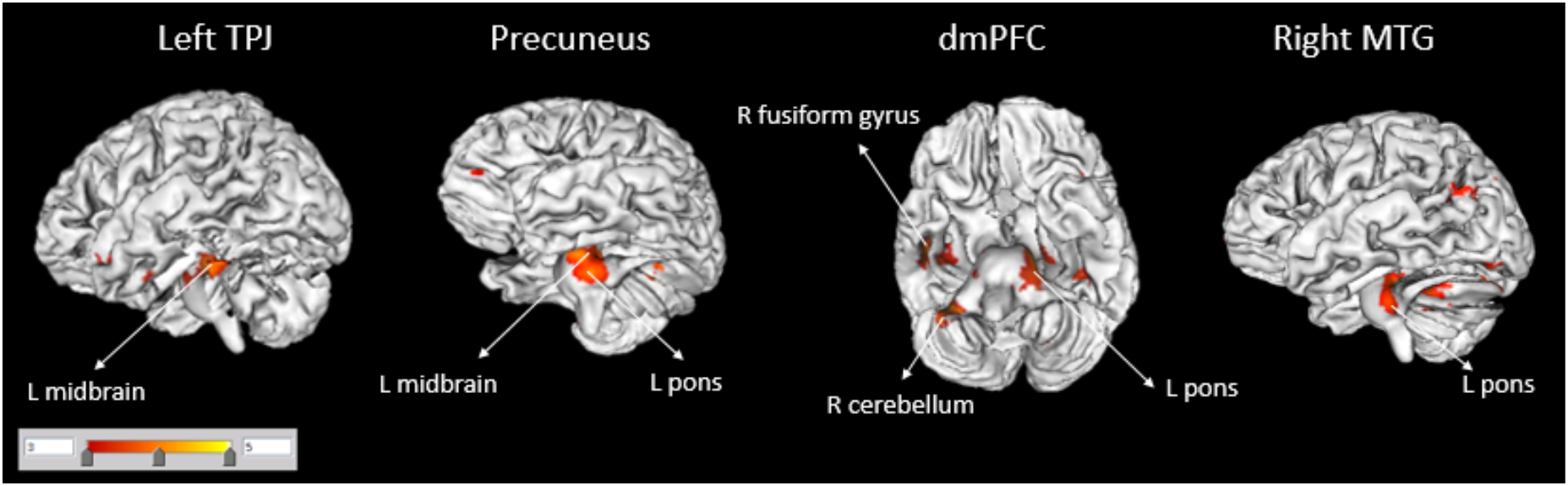
Figure shows functional connectivity of seed regions (in larger font, corrected for multiple comparisons) with other brain areas (smaller font) for the Child abuse > Unaffected comparisons contrast. There were no areas for which Unaffected comparisons > Child abuse. TPJ = temporoparietal junction; dmPFC = dorsomedial prefrontal cortex; MTG = middle temporal gyrus; R, right; L, left.

Lastly, to be confident that group differences in FC were based on paths that were existing in both groups, the spmT maps of the two groups were overlapped, revealing no significant difference (supplementary Figure 2).

**Figure 2.**
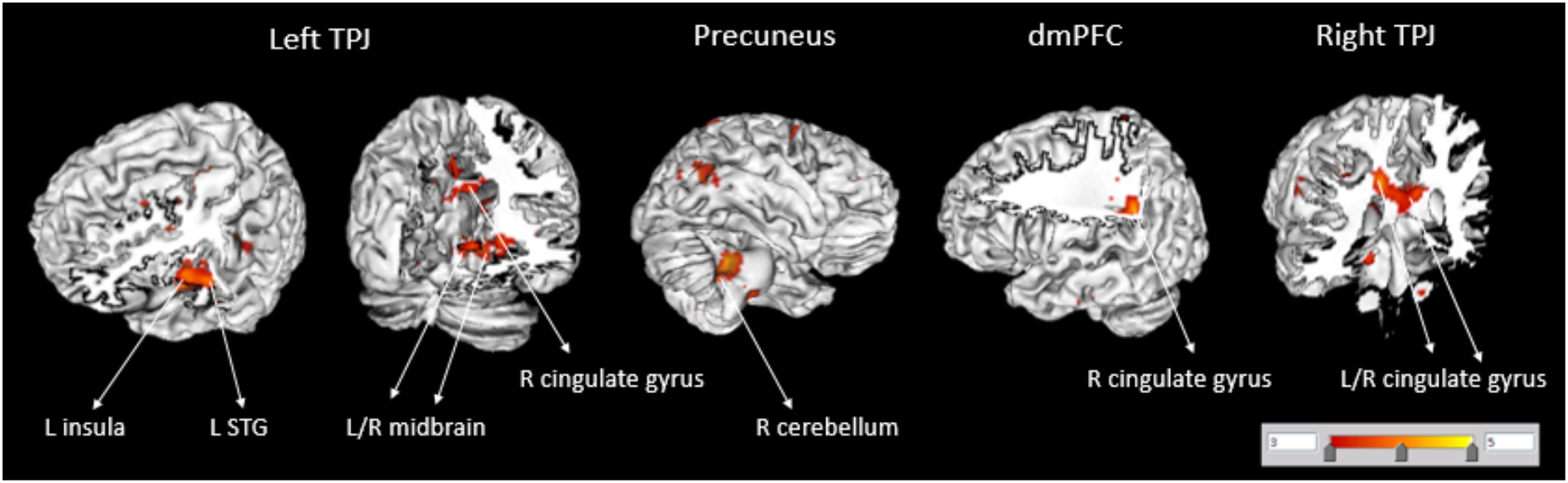
Figure shows functional connectivity of seed regions (in larger font, corrected for multiple comparisons)) with other brain areas (smaller font) for the main effect of number of abuse on the CA group. TPJ = temporo-parietal junction; dmPFC = dorsomedial prefrontal cortex; STG = superior temporal gyrus; R, right; L, left.

## Results

### Seed-based functional connectivity

ELS was associated with increased FC between the left *TPJ* and the midbrain as well as the *precuneus* and the midbrain (Table 2, Figure 1). In addition, FC was larger in previously maltreated women (vs. comparison) between the *dmPFC* seed and the anterior/posterior lobes of right cerebellum, the left pons, the right fusiform gyrus and the uncus. CA participants showed also an increased FC between the right *middle temporal gyrus* seed region and the left pons. There was no region where FC was increased for UC relative to CA.

**Table 2.**
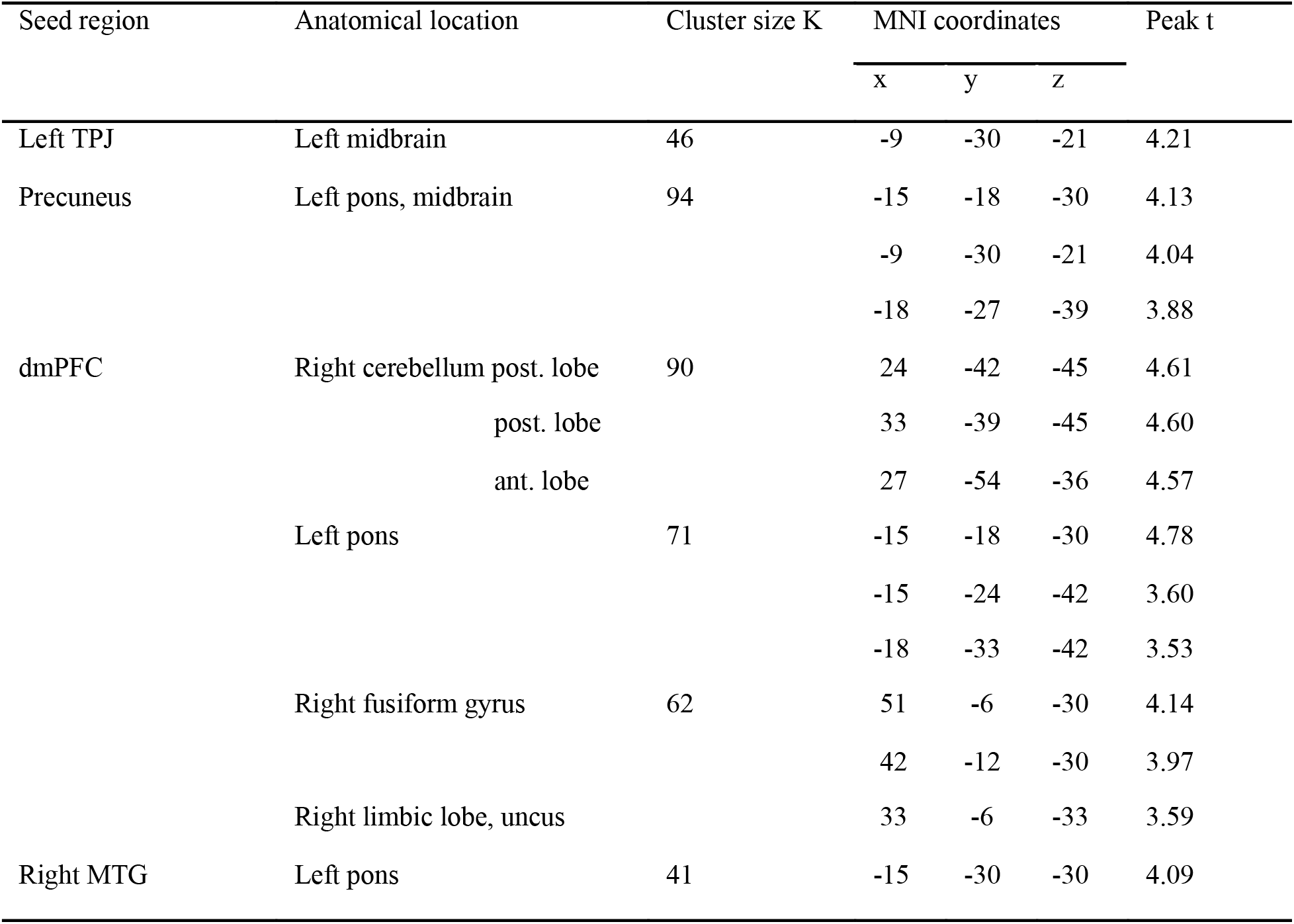
MNI coordinates and anatomical locations of the identified clusters of seed-based functional connectivity analysis. Child abuse > Unaffected comparisons

Interestingly, when the types or severity of abuse were taken into consideration, the number of types of experienced abuse revealed a significant positive FC between the left *TPJ* FC and the midbrain (Table 3, Figure 2). In other words, the more abuse a person had experienced, the stronger the FC was between the lTPJ and the midbrain. By comparison, the analysis of FC for severity of abuse revealed both positive and negative effects (Table 3) most notably a positive effect on the left *MTG* FC with right frontal regions (orbital gyrus and middle frontal gyrus) and a negative effect on the left *TPJ* FC with the left superior frontal gyrus and the left insula.

**Table 3.**
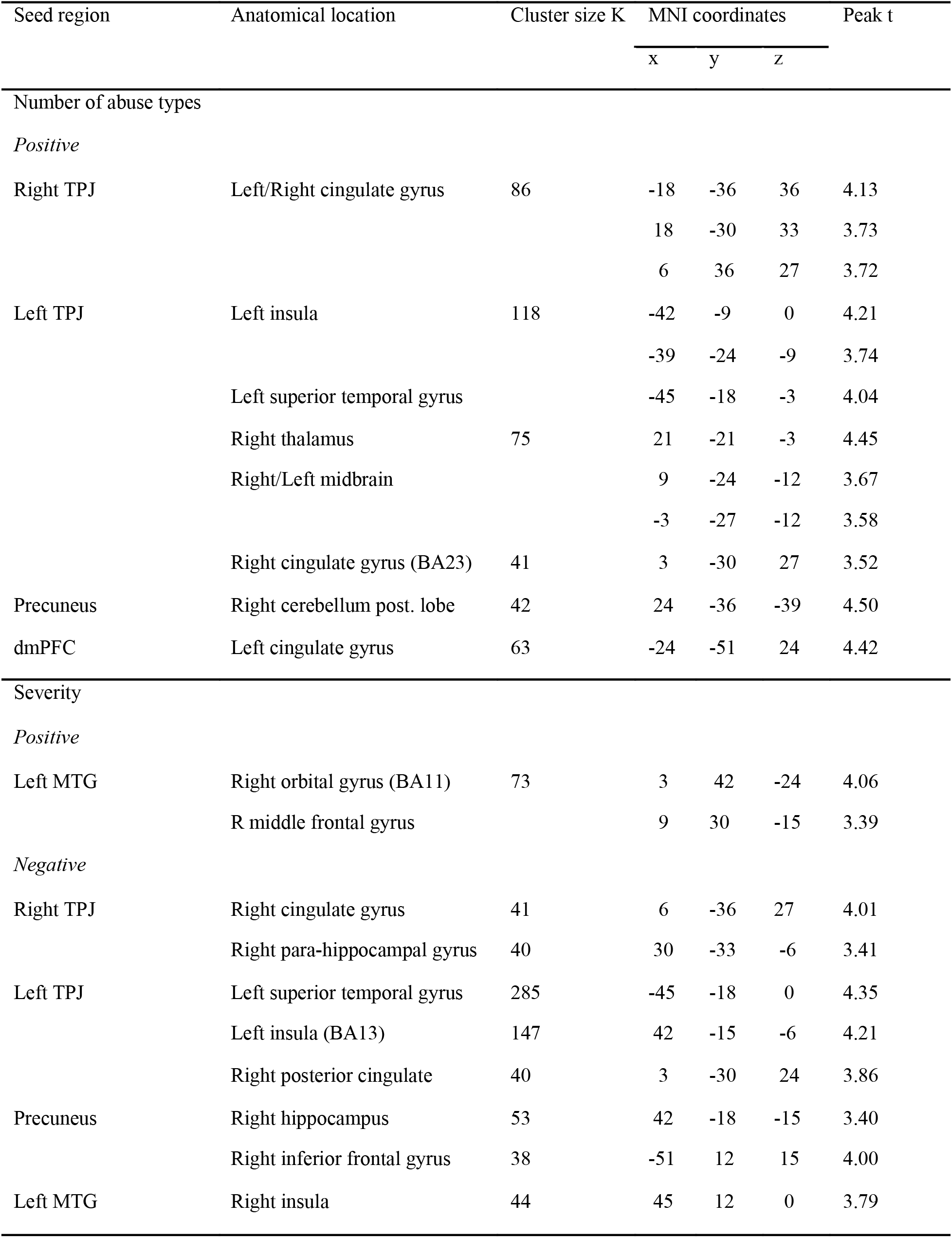
MNI coordinates and anatomical locations of the identified clusters of seed-based functional connectivity analysis for the main effect of number of abuse types and severity on the CA group

An analysis of accuracy on the ToM localizer task revealed both, a positive effect on the CA group FC and a negative effect on both groups. However, these regions were different than those relevant to the main analysis, thus not influencing the present results (cf. supplementary Table 2).

## Discussion

To our knowledge, this is the first study to investigate resting-state FC of the ToM network after ELS. Consistent with the hypothesis that an innate state of fear after abuse experience may influence social processing (12), CA women showed increased FC between most ToM seed regions and brainstem regions, whereas the dmPFC showed also connectivity to the cerebellum and the fusiform gyrus. Additional analyses revealed a positive influence of the amount of abuse experienced on the FC between the left TPJ and the midbrain.

Lanius and colleagues (11) hypothesize that traumatic experience alters the innate resting-state of the human brain, biasing it toward heightened sensitivity to threat, consistent with the hypervigilance (DSM-V) frequently reported by traumatized individuals. To date, evidence for this hypothesis has been limited as support comes from studies with small samples (12) or studies where FC for midbrain connections differed between a PTSD group without relative to those with dissociative experience but neither from controls (11). The present study focused on resting-state ToM seed regions derived from an independent localizer task, yet the brainstem emerged consistently at the whole brain level after correction for multiple comparisons, but was not one of those seed regions. FC between these brainstem/midbrain areas including the SC and LC and several important ToM regions including the TPJ, the precuneus, the right MTG and the dmPFC was increased relative to comparisons. Our data therefore support a role of brainstem regions in biasing the resting-state network after trauma. Increased connectivity with the midbrain would then indeed suggest a biasing of processing toward potential threat with the present findings extending this to a social network context. The fact that this positive FC increased with the number of abuse types experienced further cements this hypothesis. However, the precise mechanisms mediating this effect are presently poorly understood.

Despite these exciting findings, several potential explanations regarding the underlying mechanisms and neurochemistry are currently plausible that demand further inquiry. One possible explanation for the strong midbrain finding is that the midbrain is the starting point of the dopamine and serotonin pathways, which then project to wide areas of the brain including innervations to the dmPFC and the TPJ (32). Whereas the ventral tegmental area (VTA) is the source of the mesocorticolimbic dopamine pathway, the raphe nuclei are clustered along the midline throughout the brainstem, constituting the source of serotoninergic innervation (33). According to Abu-Akel (34), the integrity of the dopaminergic and serotoninergic systems is the neurochemical basis that sustains ToM ability and abnormalities in this system can account for ToM impairments. Therefore, increased FC of many ToM regions with the area from which these two important neurotransmitter pathways originate may point to a disruption in the dopaminergic-serotoninergic system that supports ToM ability in women with a history of childhood maltreatment. However, a potential alternative explanation highlights the possible complex interactions between neurochemical systems. In a recent intriguing rodent study, oxytocin release in the VTA elicited social reward and was able to influence dopaminergic neurons in this area (35) suggesting rudimentary social processes in the brainstem and an interaction between neurotransmitter systems. Yet, at this stage, such conjectures are purely speculative until more direct evidence becomes available.

Non-molecular explanations can also be found on a higher, socio-cognitive level. Recently, a greater network centrality of the precuneus in maltreated persons has been documented (36). Given its role in self-referential 1) thinking and 2) mental imagery (37), Teicher et al. suggested that the increased precuneus centrality in maltreated individuals may lead to “a heightened experience of internal emotions and cravings along with a greater tendency to think about oneself and to engage in self-centred mental imagery (p.255)” (38). Consistent with this idea, in the present data CA women relative to UC women showed increased precuneus-brainstem coupling. This suggests that women with an abuse history have an alteration in the regulation of fear responses (via the brainstem)(39) related to an increased focus on internal emotional and fearful feelings and harmful self-centred thinking.

As commonly the case with maltreated samples, findings have to be taken with caution due to a small and heterogeneous sample. To counteract this, we opted to rely on a purely female sample (in contrast to much other work, cf. (8, 9, 40, 41)) to exclude potential gender confounds. This may of course limit generalizability. A second limitation was that CM was measured retrospectively using a self-report questionnaire and is therefore sensitive to subjectivity and recall biases. Moreover, because the aim was to capture a community sample (rather than rely on a more restricted PTSD diagnosis) no clinician-administered PTSD scales (such as the CAPS) were given. However, in contrast to prior resting-state FC studies in such cohorts, we relied on an independent localizer task to ensure adequacy of the probed regions in relation to ToM function in the present sample, and also used state of the art motion correction to account for motion artefacts thus increasing sensitivity.

In summary, this study demonstrates increased FC in a theory of mind network in a community sample of women with early life maltreatment experience relative to unaffected comparisons. Particularly, increased FC consistently connected ToM regions (TPJ, dmPFC, precuneus, middle temporal gyrus) with areas of the brainstem/midbrain. The data thus strongly support prior suggestions that higher order socio-affective function may be biased already at the brainstem level in resting neural oscillations. Future work could examine whether such resting-state normalizes after therapeutic intervention. Moreover, the neurochemical role of this effect remains to be determined.

## Acknowledgements

The authors would like to warmly thank all women who participated in our research, and all organisations and groups who helped with recruitment.

---

**Supplementary Figure 1**. Figure shows functional connectivity (corrected for multiple comparisons) of seed regions (in larger font) with other brain areas (smaller font) for the Child abuse > Unaffected comparisons contrast without participants with a diagnosis of PTSD. There were no areas for which Unaffected comparisons > Child abuse. TPJ = temporo-parietal junction; dmPFC = dorsomedial prefrontal cortex; R, right; L, left.

**Supplementary Figure 2**. Figure shows functional connectivity of the dmPFC seed region with the rest of the brain for the CA and UC groups. Red = UC, green = CA, yellow = overlap. T threshold = 3.46, p = 0.001. The same pattern was displayed for the other seed regions with the same T threshold.

**Supplementary Table 1.** MNI coordinates and anatomical locations of the identified clusters of seed-based functional connectivity analysis, excluding participants with PTSD (7). Child abuse > Unaffected comparisons

| Seed region | Anatomical location | Cluster size K | MNI coordinates |  |  | Peak t |
| --- | --- | --- | --- | --- | --- | --- |
|  |  |  | x | y | z |  |
| Left TPJ | Left cerebellum ant. lobe | 51 | -12 | -36 | -18 | 4.19 |
|  | Left midbrain, pons |  | -6 | -30 | -21 | 4.13 |
|  |  |  | -15 | -18 | -30 | 3.47 |
| Precuneus | Left cerebellum ant. lobe | 47 | -12 | -33 | -24 | 3.96 |
|  | Left pons |  | -15 | -18 | -30 | 3.88 |
| dmPFC | Right cerebellum post. lobe | 125 | 33 | -39 | -45 | 4.98 |
|  | ant. lobe |  | 21 | -57 | -36 | 4.83 |
|  | post. lobe |  | 45 | -66 | -42 | 3.86 |
|  | Left pons | 76 | -12 | -21 | -27 | 4.76 |
|  | Left cerebellum |  | -18 | -30 | -45 | 3.81 |
|  | Left medulla |  | -6 | -33 | -51 | 3.36 |
|  | Right fusiform gyrus | 74 | 51 | -6 | -30 | 4.10 |
|  |  |  | 42 | -12 | -30 | 4.06 |
|  |  |  | 48 | -18 | -27 | 3.83 |

**Supplementary Table 2.** MNI coordinates and anatomical locations of the identified clusters of seed-based functional connectivity analysis for the main effect of accuracy.

| Seed region | Anatomical location | Cluster size K | MNI coordinates |  |  | Peak t |
| --- | --- | --- | --- | --- | --- | --- |
|  |  |  | x | y | z |  |
| UC |  |  |  |  |  |  |
| Negative |  |  |  |  |  |  |
| Left MTG | Left superior parietal lobule | 45 | -33 | -45 | 75 | 4.30 |
| Right MTG | Left middle frontal gyrus | 50 | -3 | 27 | -18 | 4.55 |
|  | Left postcentral gyrus | 49 | -42 | -27 | 69 | 3.98 |
| CA |  |  |  |  |  |  |
| Positive |  |  |  |  |  |  |
| dmPFC | Left cerebellum post. lobe | 41 | -48 | -66 | -30 | 4.15 |
| Negative |  |  |  |  |  |  |
| Left TPJ | Left gyrus rectus | 45 | -9 | 18 | -15 | 4.43 |

## References

1. European Union Agency for Fundamental Rights. Violence against women: an EU-wide survey. Main results report. Luxembourg: 2014.

2. Green JG, McLaughlin KA, Berglund PA, Gruber MJ, Sampson NA, Zaslavsky AM, et al. Childhood adversities and adult psychiatric disorders in the national comorbidity survey replication I: associations with first onset of DSM-IV disorders. Archives General Psychiatry. 2010;67(2):113’23. Epub 2010/02/04. doi: 67/2/113 [pii] 10.1001/archgenpsychiatry.2009.186 [doi]. PubMed PMID: 20124111; PubMed Central PMCID: PMC2822662.

3. Taillieu TL, Brownridge DA, Sareen J, Afifi TO. Childhood emotional maltreatment and mental disorders: Results from a nationally representative adult sample from the United States. Child abuse & neglect. 2016;59:1–12. Epub 2016/08/05. doi: 10.1016/j.chiabu.2016.07.005. PubMed PMID: 27490515.

4. Luke N, Banerjee R. Differentiated associations between childhood maltreatment experiences and social understanding: A meta-analysis and systematic review. Developmental Review. 2013;33(1):1–28. doi: 10.1016/j.dr.2012.10.001.

5. Gallagher HL, Frith CD. Functional imaging of ‘theory of mind’. Trends in cognitive sciences. 2003;7(2):77–83. Epub 2003/02/14. PubMed PMID: 12584026.

6. Schurz M, Radua J, Aichhorn M, Richlan F, Perner J. Fractionating theory of mind: a meta-analysis of functional brain imaging studies. Neurosci Biobehav R. 2014;42:9–34. Epub 2014/02/04. doi: 10.1016/j.neubiorev.2014.01.009. PubMed PMID: 24486722.

7. van Veluw SJ, Chance SA. Differentiating between self and others: an ALE metaanalysis of fMRI studies of self-recognition and theory of mind. Brain imaging and behavior. 2014;8(1):24–38. Epub 2014/02/19. doi: 10.1007/s11682-013-9266-8 [doi]. PubMed PMID: 24535033.

8. Thomason ME, Marusak HA, Tocco MA, Vila AM, McGarragle O, Rosenberg DR. Altered amygdala connectivity in urban youth exposed to trauma. Social cognitive and affective neuroscience. 2015;10(11):1460–8. Epub 2015/04/04. doi: 10.1093/scan/nsv030. PubMed PMID: 25836993; PubMed Central PMCID: PMCPMC4631140.

9. Philip NS, Sweet LH, Tyrka AR, Price LH, Bloom RF, Carpenter LL. Decreased default network connectivity is associated with early life stress in medication-free healthy adults. European neuropsychopharmacology: the journal of the European College of Neuropsychopharmacology. 2013;23(1):24–32. Epub 2012/11/13. doi: 10.1016/j.euroneuro.2012.10.008. PubMed PMID: 23141153; PubMed Central PMCID: PMCPMC3581700.

10. van der Werff SJ, Pannekoek JN, Veer IM, van Tol MJ, Aleman A, Veltman DJ, et al. Resting-state functional connectivity in adults with childhood emotional maltreatment. Psychological medicine. 2013;43(9):1825–36. Epub 2012/12/21. doi: 10.1017/s0033291712002942. PubMed PMID: 23254143.

11. Olive I, Densmore M, Harricharan S, Theberge J, McKinnon MC, Lanius R. Superior colliculus resting state networks in post-traumatic stress disorder and its dissociative subtype. Human brain mapping. 2018;39(1):563–74. Epub 2017/11/15. doi: 10.1002/hbm.23865. PubMed PMID: 29134717.

12. Steuwe C, Daniels JK, Frewen PA, Densmore M, Theberge J, Lanius RA. Effect of direct eye contact in women with PTSD related to interpersonal trauma: Psychophysiological interaction analysis of connectivity of an innate alarm system. Psychiatry research. 2015;232(2):162–7. Epub 2015/04/12. doi: 10.1016/j.pscychresns.2015.02.010. PubMed PMID: 25862529.

13. Van der Kolk BA. The body keeps the score. New York: Penguin Books; 2015.

14. Steuwe C, Daniels JK, Frewen PA, Densmore M, Pannasch S, Beblo T, et al. Effect of direct eye contact in PTSD related to interpersonal trauma: an fMRI study of activation of an innate alarm system. Social cognitive and affective neuroscience. 2014;9(1):88–97. Epub 2012/09/15. doi: 10.1093/scan/nss105. PubMed PMID: 22977200; PubMed Central PMCID: PMCPMC3871730.

15. Schmidt U, Willmund GD, Holsboer F, Wotjak CT, Gallinat J, Kowalski JT, et al. Searching for non-genetic molecular and imaging PTSD risk and resilience markers: Systematic review of literature and design of the German Armed Forces PTSD biomarker study. Psychoneuroendocrinology. 2015;51:444–58.

16. Dodell-Feder D, Koster-Hale J, Bedny M, Saxe R. fMRI item analysis in a theory of mind task. NeuroImage. 2011;55(2):705–12. Epub 2010/12/25. doi: 10.1016/j.neuroimage.2010.12.040. PubMed PMID: 21182967.

17. Goodman LA, Corcoran C, Turner K, Yuan N, Green BL. Assessing traumatic event exposure: general issues and preliminary findings for the Stressful Life Events Screening Questionnaire. Journal of traumatic stress. 1998;11(3):521–42. Epub 1998/08/05. doi: 10.1023/a:1024456713321. PubMed PMID: 9690191.

18. Bernstein DP, Fink L. Childhood Trauma Questionnaire: A retrospective self-report manual. San Antonio, TX: The Psychological Corporation; 1998.

19. Davis MH. Measuring individual differences in empathy: Evidence for a multidimensional approach. Journal of Personality and Social Psychology. 1983;44(1): 113–26. doi: http://dx.doi.org/10.1037/0022-3514.44.L113

20. De Corte K, Buysse A, Verhofstadt LL, Roeyers H, Ponnet K, Davis MH. Measuring Empathic Tendencies: Reliability And Validity of the Dutch Version of the Interpersonal Reactivity Index Psychologica Belgica 2007;47(4):235–60. doi: http://doi.org/10.5334/pb-47-4-235

21. Beck AT, Steer RA, Brown GK. Manual for the Beck Depression Inventory-II. Second ed. San Antonio; TX: Psychological Corporation; 1996.

22. Spielberger CD, Gorsuch RL, Lushene R, Vagg PR, Jacobs GA. Manual for the State-Trait Anxiety Inventory. Palo Alto, CA, US: Consulting Psychologists Press; 1983.

23. Van der Ploeg HM, Defares PB, Spielberger CD. Handleiding bij de Zelf-Beoordelings Vragenlijst ZBV. Een nederlandstalige bewerking van de Spielberger State-Trait Anxity Inventory [Manual of the Dutch State-Trait Anxiety Inventory]. Lisse, Netherlands: Swets & Zeitlinger; 1980

24. Bernstein EM, Putnam FW. Development, reliability, and validity of a dissociation scale. The Journal of nervous and mental disease. 1986;174(12):727–35. Epub 1986/12/01. PubMed PMID: 3783140.

25. Ensink BJ, van Otterloo D. A validation of the Dissociative Experiences Scale in the Netherlands. Dissociation. 1989;2(4):221–23.

26. Wagnild GM, Young HM. Development and psychometric evaluation of the Resilience Scale. J Nurs Meas. 1993;1(2):165–78. Epub 1993/01/01. PubMed PMID: 7850498.

27. Overbeek T, Schruers K, Griez E. Mini International Neuropsychiatric Interview: Nederlandse Versie 5.0.0. Universiteit van Maastricht, Maastricht, NE 1999.

28. Sheehan DV, Lecrubier Y, Harnett-Sheehan K, Janavas J, Weiller E, Bonara LI, et al. Reliability and Validity of the M.I.N.I. International Neuropsychiatric Interview (M.I.N.I.): According to the SCID-P. European Psychiatry. 1997;12:232–41.

29. Cordes D, Haughton VM, Arfanakis K, Carew JD, Turski PA, Moritz CH, et al. Frequencies contributing to functional connectivity in the cerebral cortex in ‘resting-state’ data. AJNR Am J Neuroradiol. 2001;22(7):1326–33. Epub 2001/08/11. PubMed PMID: 11498421.

30. Power JD, Barnes KA, Snyder AZ, Schlaggar BL, Petersen SE. Spurious but systematic correlations in functional connectivity MRI networks arise from subject motion. NeuroImage. 2012;59(3):2142–54. Epub 2011/10/25. doi: 10.1016/j.neuroimage.2011.10.018. PubMed PMID: 22019881; PubMed Central PMCID: PMCPMC3254728.

31. Power JD, Mitra A, Laumann TO, Snyder AZ, Schlaggar BL, Petersen SE. Methods to detect, characterize, and remove motion artifact in resting state fMRI. NeuroImage. 2014;84:320–41. Epub 2013/09/03. doi: 10.1016/j.neuroimage.2013.08.048. PubMed PMID: 23994314; PubMed Central PMCID: PMCPMC3849338.

32. Abu-Akel A, Shamay-Tsoory S. Neuroanatomical and neurochemical bases of theory of mind. Neuropsychologia. 2011;49(11):2971–84. Epub 2011/08/02. doi: 10.1016/j.neuropsychologia.2011.07.012. PubMed PMID: 21803062.

33. Venkatraman A, Edlow BL, Immordino-Yang MH. The Brainstem in Emotion: A Review. Frontiers in neuroanatomy. 2017;11:15. Epub 2017/03/25. doi: 10.3389/fnana.2017.00015. PubMed PMID: 28337130; PubMed Central PMCID: PMCPMC5343067.

34. Abu-Akel A. The neurochemical hypothesis of ‘theory of mind’. Med Hypotheses. 2003;60(3):382–6. Epub 2003/02/13. PubMed PMID: 12581615.

35. Hung LW, Neuner S, Polepalli JS, Beier KT, Wright M, Walsh JJ, et al. Gating of social reward by oxytocin in the ventral tegmental area. Science. 2017;357(6358):1406–11.

36. Teicher MH, Anderson CM, Polcari A. Childhood maltreatment: Altered network centrality of cingulate, precuneus, temporal pole and insula. Biological Psychiatry. 2014;76:297–305.

37. Cavanna AE, Trimble MR. The precuneus: a review of its functional anatomy and behavioural correlates. Brain: a journal of neurology. 2006;129(Pt 3):564–83. PubMed PMID: 16399806.

38. Teicher MH, Samson JA. Annual Research Review: Enduring neurobiological effects of childhood abuse and neglect. Journal of child psychology and psychiatry, and allied disciplines. 2016;57(3):241–66. Epub 2016/02/03. doi: 10.1111/jcpp.12507. PubMed PMID: 26831814; PubMed Central PMCID: PMCPMC4760853.

39. Adolphs R. The Biology of Fear. Curr Biol. 2013;23(2):R79–93. doi: 10.1016/j.cub.2012.11.055 [doi]. PubMed PMID: 23347946.

40. Lu S, Gao W, Wei Z, Wang D, Hu S, Huang M, et al. Intrinsic brain abnormalities in young healthy adults with childhood trauma: A resting-state functional magnetic resonance imaging study of regional homogeneity and functional connectivity. Aust N Z J Psychiatry. 2017;51(6):614–23. Epub 2016/10/04. doi: 10.1177/0004867416671415. PubMed PMID: 27694638.

41. Herringa RJ, Birn RM, Ruttle PL, Burghy CA, Stodola DE, Davidson RJ, et al. Childhood maltreatment is associated with altered fear circuitry and increased internalizing symptoms by late adolescence. Proceedings of the National Academy of Sciences of the United States of America. 2013;110(47):19119–24. Epub 2013/11/06. doi: 10.1073/pnas.1310766110. PubMed PMID: 24191026; PubMed Central PMCID: PMCPMC3839755.

